# TEP-1, a glial thioester protein is required for cilia organization and intraflagellar transport in ensheathed sensory neurons

**DOI:** 10.1101/2025.10.23.684198

**Authors:** Yumiko Oshima, Yaminisree Nagidi, Maya E. Moorthy, Jonathon Heier, Joanne A. Matsubara, Jeff Hardin, Martin Flajnik, Bruce E. Vogel

**Affiliations:** Center for Biomedical Engineering and Technology and Department of Physiology, University of Maryland School of Medicine, University of Maryland, Baltimore, MD 21201; Department of Integrative Biology, University of Wisconsin Madison, Madison, WI 53706; Department of Ophthalmology and Visual Sciences, University of British Columbia, Vancouver, BC V5Z 3N9, Canada; Department of Microbiology and Immunology, University of Maryland School of Medicine, University of Maryland, Baltimore, MD 21201

## Abstract

Age-related macular degeneration (AMD), the leading cause of blindness in the elderly, is characterized by progressive degeneration of retinal photoreceptors. Traditional disease models suggest that defective repression of thioester protein C3 activity by complement factor H (CFH) is a major contributor to pathogenesis in AMD and a related disease, early-onset drusen maculopathy (EODM). Our previous study identified novel functions for human CFH and *C. elegans* CFH-1 in the maintenance of inversin compartment integrity in photoreceptors and mechanosensory neurons, indicating that CFH has a novel, evolutionarily conserved role in cilia compartment organization that is distinct from its established function in alternative complement pathway regulation. Here, we investigate the *C. elegans* thioester protein TEP-1, an ancestral relative of C3 and other members of the AMCOM family (C4, C5, CD109, and alpha-2-macroglobulin). TEP-1 localizes to select glial cell surfaces and regulates inversin compartment organization and intraflagellar transport (IFT) within the cilia of ensheathed sensory neurons. In addition to revealing a novel role for an AMCOM family member in sensory neuron structure and protein transport, the localization of C3 and CFH on human photoreceptors provides support for non-canonical models of AMD and EODM pathogenesis in which defects in cilia structure and protein transport contribute directly to the progressive photoreceptor dysfunction that characterizes these diseases.

## Introduction

Age-related macular degeneration (AMD) is a leading cause of blindness among the elderly, affecting 11% of adults over the age of 85, although the mechanisms of disease initiation and progression are not fully understood. Defective regulation of the alternative complement pathway, part of the innate immune system’s natural defense against infections, was implicated as a key contributor to AMD progression by the discovery of amino acid substitutions (Y402H and R1210C) in complement factor H (CFH) as AMD risk factors (1–6). CFH is a secreted regulatory protein that promotes decay of the C3bBb convertase that activates thioester protein C3 and acts as a cofactor for the factor I protease in processing C3b to its inactive form (7). Current models for AMD suggest that the accumulation of drusen, extracellular deposits located between the retinal pigment epithelium (RPE) and Bruch’s membrane, coupled with defective regulation of C3 by CFH, result in inflammation and cytolysis that disrupt the function of RPE cells and their glial-like interactions with photoreceptors (8,9). Genetic studies of patients with early onset drusen maculopathy (EODM), a disease related to AMD, identified CFH sequence variants in 30.3% of patient samples. Most of these mutations were found in regions where CFH binds C3 (10,11). While these findings are consistent with inflammation-based models of AMD and EODM pathogenesis, CFH has non-canonical functions in monocyte migration, lipid distribution, and cilia compartment organization that may influence AMD pathogenesis (12–14).

Although *C. elegans* does not have an elaborate visual system with specialized photoreceptors, 60 ciliated sensory neurons that are ensheathed by glia detect chemical, thermal, and mechanical stimuli (15–17). Like their vertebrate counterparts, nematode cilia are microtubule-based cell protrusions with evolutionarily conserved protein compositions and compartmental organizations (16,18). In between the proximal transition zone (TZ) and distal segment (DS) is the middle segment (MS) that in many cilia subtypes contains a subdomain called the “inversin compartment,” defined by the location of the inversin protein [also called NPHP-2 (19, 20)]. In a previous study we demonstrated that loss-of-function mutations in the *C. elegans* CFH homolog, CFH-1, result in progressive defects in inversin compartment integrity and cilia function in the sensory neurons of aging animals (14). Similar defects in inversin/NPHP-2 distribution were also detected in human photoreceptors from individuals homozygous for the AMD high-risk CFH Y402H variant, suggesting that cilia structural defects may contribute to the progressive photoreceptor dysfunction observed in AMD patients with CFH loss-of-function mutations (14).

C3, the main regulatory target of CFH, is a member of the vertebrate α2-macroglobulin/complement (AMCOM) protein family, that includes C4, C5, and CD109 (21). Although C3 activity is thought to have a detrimental effect on photoreceptor health in the absence of CFH-mediated inhibition, knockout mouse studies indicate that C3 actually has a protective effect on photoreceptor health that is independent of its interaction with CFH (22). Together with recent discoveries of novel, non-canonical CFH functions, this suggests that CFH and C3 have poorly understood functions outside of their established roles in the alternative complement pathway that influence photoreceptor health (12–14, 22). Here, we investigate the distribution and function of TEP-1, a *C. elegans* ancestral relative of the AMCOM family, and find that TEP-1 localizes to the cell surfaces of select glial cells that ensheath the cilia of chemosensory and mechanosensory neurons (17). Our data suggest that TEP-1, like CFH-1, is required for inversin/NPHP-2 organization within its namesake compartment and the regulation of intraflagellar transport (IFT) rates in ensheathed sensory neuron cilia. Together with recent demonstrations of CFH and CFH-1 function in cilia structure and IFT (14,23), this study supports a model for AMD and EODM pathogenesis that involves contributions from primary defects in photoreceptor cilia structure and protein transport.

## Results

### Identification of TEP-1, the founding member of the AMCOM family

The vertebrate AMCOM family of thioester proteins includes complement components C3, C4, C5, CD109, and α2-macroglobulin (21). Each family member has several conserved sequence motifs, including a furin cleavage site and, with the exception of C5, the thioester linkage consensus sequence GCGEQ that is capable of forming a covalent bond with substrates through thioester transacylation. TEP-1 was identified as a *C. elegans* AMCOM family ancestral relative (21) (Fig. S1) and has multiple mRNA alternate splice forms (https://wormbase.org/species/c_elegans/gene/WBGene00013969#0-9fdc41beg365hi-10). At least one splice form contains a C-terminal GPI anchor sequence found only in CD109 and not in other vertebrate AMCOM members (21). The furin cleavage site location and the absence of a GPI anchor sequence in a second TEP-1 splice variant make it structurally similar to C3 and C4 although C3, C4, and C5 contain anaphylatoxin and netrin domains not found in α2-macroglobulin, CD109 or TEP-1 (Fig. S1A).

### TEP-1 is expressed and localizes to the distal ends of a subset of sensory neuron glial cells

To investigate *tep-1* expression, a transcriptional reporter construct containing approximately 3.0 kb of *tep-1* 5’ regulatory sequence immediately upstream of GFP was introduced into wild-type (WT) animals. *tep-1p*::GFP is expressed in a subset of sensory neuron glial cells including CEP sheath (CEPsh) and phasmid socket glia (Phso) based on cell position, shape, and overlap with cellular expression markers including *ptr-10p*::myrRFP (Fig. S1B)(24, 25). Expression by these sensory neuron glial cells was notable because of the potential overlap of TEP-1 with CFH-1, previously identified on the surface of CEP neurons and phasmid sheath glia (Phsh) (14), and because a central element of the alternative complement pathway and AMD and EODM pathogenesis is thought to be a defect in CFH-mediated regulation of C3, a TEP-1-related thioester protein.

To determine TEP-1 protein localization, mNeon Green (mNG) was inserted in-frame into endogenous *C. elegans* TEP-1 coding sequence between the signal peptide cleavage site and the first conserved MG2 domain (Fig. S1A) using CRISPR-Cas-9 to create an mNG::TEP-1 fusion protein. Subcellular distribution of mNG::TEP-1 was observed at the distal ends of CEPsh and amphid socket (Amso) glial cells based on position and partial overlap in distribution with multiple neuronal and glial markers including *ptr-10p*::myrRFP and *grl-2p*::mApple (24,25) (Fig. 1). mNG::TEP-1 concentrates at the distal ends of these tube-shaped glial cells where it overlaps VAB-9, a member of the claudin family of tetraspanin proteins that localizes at *C. elegans* adherens junctions (Fig. 1B) (26). In glial cells, adherens junctions connect sheath to socket glia and dendrites, and socket glia to hypodermis (27). TEP-1 also localizes at the distal ends of Phso glial cells and extends proximally along what appears to be Phso and Amso inner surfaces, adjacent to VAB-9 (Fig. 1B). mNG::TEP-1 distribution also overlaps VAB-9 in seam and vulval epithelia, and is found in the interstitial fluid of the uterus and pseudocoelomic cavity. (Fig. S1C).

**Figure 1.**
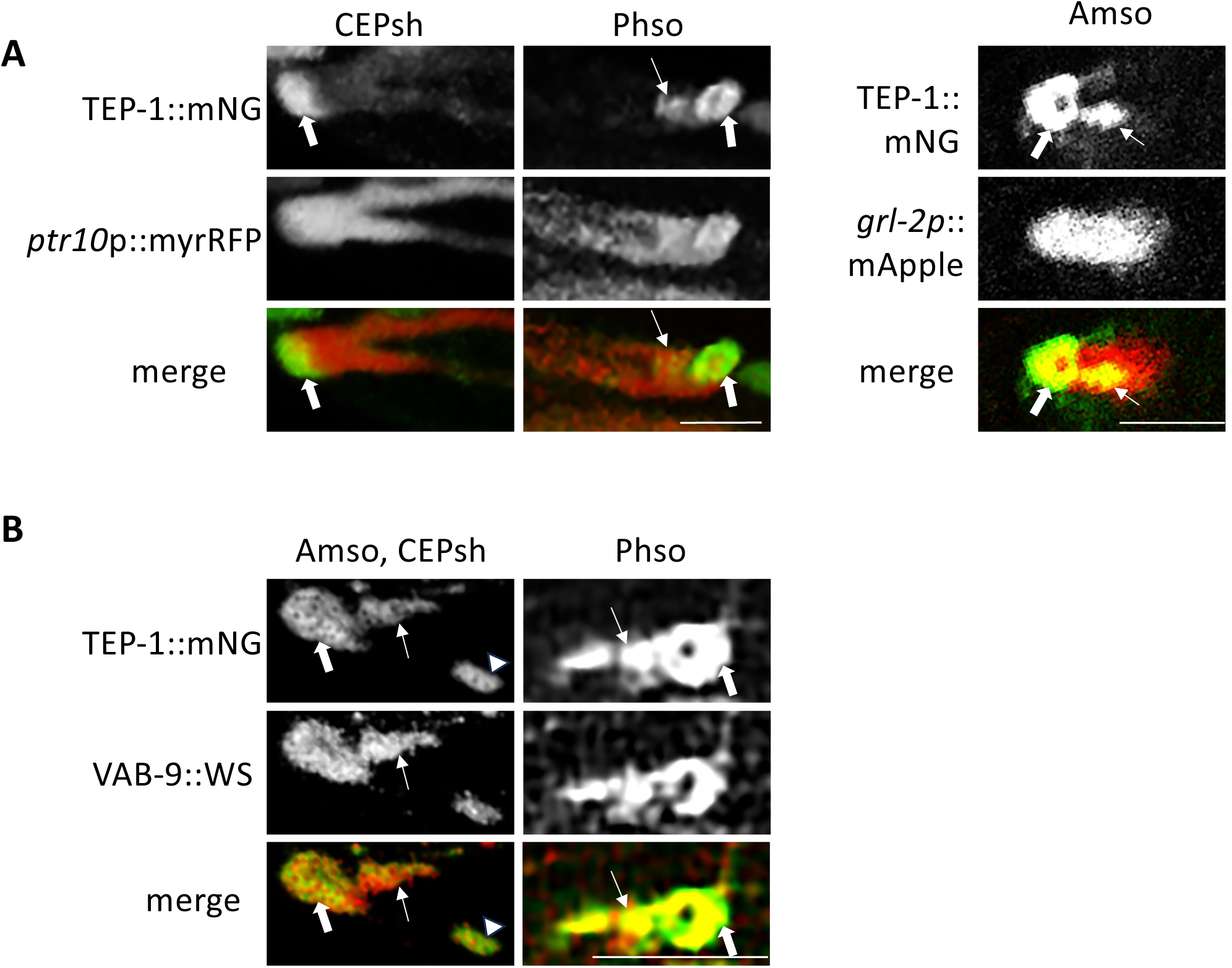
*TEP-1::mNG localization on* sensory neurons glial cells. A, TEP-1::mNG translational fusion protein (thick arrrows) is detected at the distal ends of 4 CEP sheath (CEPsh), 2 Phasmid socket (Phso), and 2 Amphid socket (Amso) tube-shaped glial cells indicated by overlap with *ptr-10p::mRFP* or *grl-2::mApple* (red). TEP-1::mNG also extends proximally on Amso and Phso (thin arrows). Anterior is oriented left. Scale bars are 5 μm. B, Confocal image showing TEP-1::mNG and VAB-9::wrmScarlet translational fusion proteins overlap at distal ends of Amso (thick arrows), CEPsh (arrowheads), and Phso (thick arrows) glial cells. TEP-1::mNG and VAB-9::wrmScarlet also extend proximally form the distal end of the Amso and Phso tube-shaped glial cells (thin arrows). Scale bars are 5μm. Anterior is oriented left.

### TEP-1 encircles sensory neuron cilia

The distal ends of socket and sheath glia encircle cilia that extend from sensory neuron dendrites. To evaluate glial TEP-1 localization with respect to sensory neuron cilia, mNG::TEP-1 was imaged in tandem with dynein light-intermediate chain protein XBX-1::tagRFP. TEP-1 CEPsh glia encircles XBX-1 on the cilia axoneme, approximately 2 μm from the base and 2.5-3 μm from the distal end of CEP sensory neuron cilia (Fig. 2A, 2C). On phasmid sensilla, the mNG::TEP-1 torus at the distal end of phasmid socket glia encircles XBX-1::tagRFP at the distal end of the phasmid axoneme and forms an elongated bowl that extends approximately 3 μm proximally along the inner surface of the phasmid socket cell, adjacent to the phasmid axoneme (Fig. 2A, 2C).

**Figure 2.**
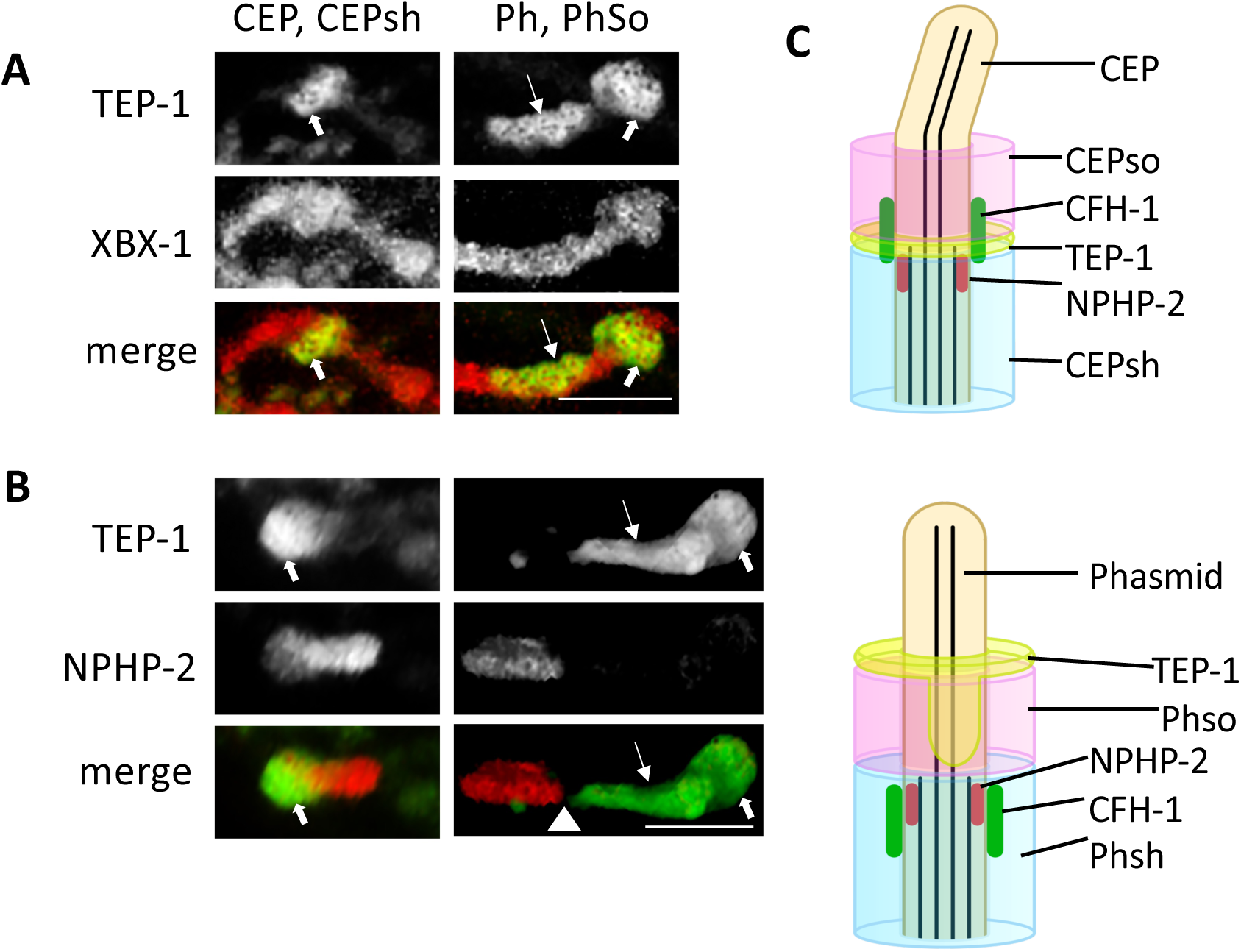
Glial TEP-1 encircles CEP and Phasmid sensory neuron cilia. *A, TEP-1::mNG encircles XBX-1 dynein subunit on CEP and Phasmid (Ph) sensory neuron cilia.* TEP-1 at the distal end of CEPsh and Phso glia (thick arrows) and along the phasmid distal segment (thin arrows) encircles XBX-1 in CEP and Phasmid cilia. Anterior oriented left. Scale bar is 2.5μm B, TEP-1::mNG is located at the distal ends of the CEP and phasmid inversin compartments. The location of TEP-1 at the distal end of CEPsh and PHso glia (thick arrows) and along the phasmid distal segment (thin arrows) is shown. The gap between Inversin/NPHP-2 and TEP-1 is indicated by the arrowhead. Anterior oriented left. Scale bar is 2.5μm *C,* Schematic diagram showing arrangement of inversin/NPHP-2, CFH-1 and TEP-1 in relationship to socket (pink) and sheath (blue) glial cells on CEP (top) and phasmid sensilla.

Cilia are divided into discrete compartments that include a proximal transition zone (TZ), middle segment (MS), and distal segment (DS). To refine TEP-1 localization with respect to these cilia compartments, mNG::TEP-1 was imaged in tandem with inversin/NPHP-2::mCh, the protein that defines the ‘inversin’ compartment that is located within cilia middle segments. On CEPsh glia, mNG::TEP-1 flanks the distal end of inversin/NPHP-2 fibrils in CEP sensory neuron cilia with little or no detectable gap between the two proteins (Fig. 2B, 2C). mNG::TEP-1 on Phso extends nearly to the inversin/NPHP-2 fibrils of phasmid cilia, with a detectable gap of approximately 0.5μm between the two proteins (Fig. 2B, 2C).

### Glial expression of TEP-1 maintains inversin compartment organization in adjacent sensory neurons

Previous work has shown that CFH-1 is localized on CEP sensory neuron cilia, where it prevents ectopic accumulation of inversin/NPHP-2 in the distal cilia compartment of aging animals (14). Based on this CFH-1 function, the established regulatory interaction between CFH and the thioester protein C3 in vertebrates, and TEP-1 localization relative to inversin/NPHP-2 (Fig. 2B, 2C), the distribution of CFH-1 and NPHP-2 was investigated in *tep-1(em18)* and *tep-1(ok2874)* null mutant adults that have an 8bp frameshift deletion immediately after TEP-1 signal peptide coding sequence and a 360bp deletion, respectively. In CEP neurons, there is a statistically significant increase in the size of the inversin compartment in day 4 *tep-1* mutant adults (Fig. 3). Similarly, in phasmid neurons, there is also a statistically significant increase in the length and the concentration of inversin/NPHP-2 in *tep-1* mutants (Fig. 3A). The distribution of CFH-1 on CEP neurons is mildly affected in some *tep-1(ok2874)* mutants, although this effect is not statistically significant and minor when compared to the pronounced defects in CFH-1 distribution observed previously in syndecan/SDN-1 mutants (Fig. 3A) (14).

**Figure 3.**
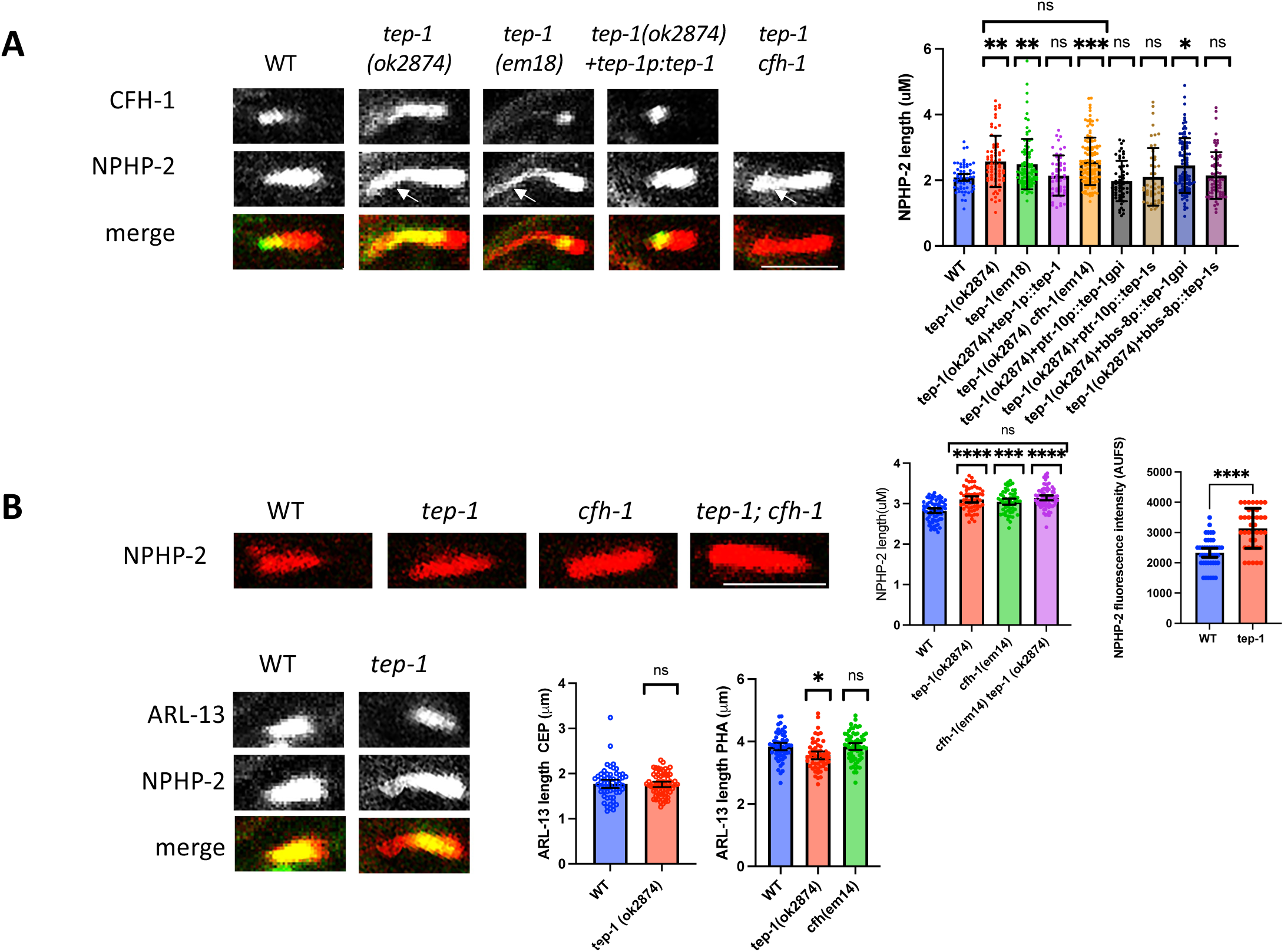
TEP-1 prevents ectopic accumulation of inversin/NPHP-2 in distal cilia compartments. A, Comparison of CFH-1::GFP and inversin/NPHP-2::mCherry localization in CEP neurons of WT, *tep-1(ok2874), tep-1(em18), tep-1(ok2874)* containing *tep-1p::tep-1* transgene, and *tep-1(ok2874);cfh-1(em14)* day 4 adult animals. Note that inversin/NPHP-2 is restricted in length in WT but extends into cilia distal segment in *tep-1* mutant animals (arrows). Right, scatter plots of inversin/NPHP-2 length in CEP neurons of day 4 adult animals. Scale bars are 2.5μm Error bars indicate 95% confidence intervals. Significance is indicated by brackets with asterisks as follows: *P<.1; ** P<.01; ***P<.001; ns indicates no significant difference from WT. B, (Top), Comparison of inversin/NPHP-2::mCherry localization in the phasmid neurons of WT, *tep-1, cfh-1* and *tep-1;cfh-1* mutant animals. Right, scatter plots of inversin/NPHP-2 length and fluorescence intensity in phasmid neurons of day 4 adult animals. Scale bars are 2.5μm. Significance is indicated by brackets with asterisks as follows: ***P<.001; ****P<.0001; ns indicates no significant difference from WT. Anterior is oriented left in all images. Bottom, ARL-13 lengths in CEP neurons of WT and tep-1 mutant animals (top) and ARL-13 length in the phasmids of WT, *tep-1 and cfh-1* mutant animals. Scale bars are 2.5μm. Right, scatter plots of ARL-13 length in CEP (top) and phasmid neurons of day 4 adult animals. Error bars indicate 95% confidence intervals. Significance is indicated by brackets with asterisks as follows: *P<.1; ns indicates no significant difference from WT.

A transgenic construct containing tep-1 5’ and 3’ regulatory sequences flanking tep-1 coding sequence successfully rescued the mutant phenotype (Fig. 3A), demonstrating that the mislocalization of inversin/NPHP-2 observed in *tep-1* mutant animals is due to *tep-1* loss of function rather than an unrelated mutation elsewhere in the genome. In addition, 4 different transgenic constructs were used to determine whether the structure of the inversin compartment in CEP neurons depends on cell autonomous neuronal or nonautonomous glial expression and whether membrane-anchored or secreted forms of TEP-1 are necessary for its function in inversin compartment organization. A *tep-1* transgenic construct containing a predominantly glial *ptr-10* promoter and GPI anchor sequences rescued the mutant phenotype while a neuronal *bbs-8* promoter construct containing the GPI anchor did not, suggesting that cell-nonautonomous glial expression is critical for TEP-1 function. In contrast, non-anchored constructs containing either *ptr-10* or *bbs-8* regulatory sequences both rescued the mutant phenotype (Fig. 3A). Together, these data suggest that TEP-1 functions in the organization of the neuronal inversin compartment when it is attached to glial cell surfaces or secreted into the space between glial cells and neurons, but not when it is attached to neuronal cell surfaces.

### TEP-1 and CFH-1 functions have partial overlap

In contrast to the inversin/NPHP-2 phenotype, there was little or no detectable difference in ARL-13::mNG distribution between WT and *tep-1* mutant animals, suggesting that the absence of TEP-1 protein has a specific effect on inversin/NPHP-2 and does not cause a general defect in cilia structure and organization in CEP or phasmid neurons (Fig. 3B). This is consistent with the inversin/NPHP-2 specific phenotype of *cfh-1* mutants seen previously (14) and suggests potential overlap in *cfh-1* and *tep-1* function.

To examine potential interaction between CFH-1 and TEP-1, a *cfh-1(em14); tep-1(ok2847)* double null mutant was tested for enhancement of the inversin/NPHP-2 phenotype in CEP and phasmid neuron cilia. A small but statistically insignificant increase was found in the size of the inversin compartment in the CEP and phasmid neurons of *cfh-1 (em14); tep-1 (ok2874)* double mutants when compared to *cfh-1(em14)* and *tep-1(ok2874)* single mutants. (Fig. 3).

### Tep-1 mutant animals have defects in intraflagellar transport (IFT)

Cilia assembly, maintenance and function are dependent on the bidirectional movement of intraflagellar transport (IFT) trains along the microtubule based cilia axoneme. IFT trains are polymers of IFT-A and IFT-B protein complexes that are transported together with IFT cargo by kinesin and dynein motor proteins (28). Based on the disruption of inversin/NPHP-2 in *tep-1* mutants, the observation that another component of the NPHP module, NPHP-4, negatively regulates IFT rates (29), and IFT defects in cfh-1 mutant animals (23), we examined whether IFT rates were affected in *C. elegans tep-1* mutant animals. Examination of IFT-A component CHE-11/IFT140, IFT-B1 component OSM-5/IFT88 IFT, and dynein heavy chain CHE-3 revealed slight, but statistically significant increases in the rate of anterograde transport in *tep-1(ok2874)* mutant animals (Fig. S3). Similarly, examination of dynein light-intermediate chain XBX-1 revealed increases in the rate of anterograde and retrograde transport in *tep-1(ok2874)* mutant animals when compared to WT (Fig. 4). In striking contrast to all other IFT train components examined, IFTB1 component, IFT52/OSM-6, had significantly slower rates of anterograde and retrograde transport in *tep-1(ok2874)* mutant animals. The data suggest distinct effects of TEP-1 on IFT52/OSM-6 and IFT88/OSM-5 rates even though both are subunits of the IFTB1 complex (28). This is similar to the effect of CFH-1 on these IFT train components (23) and provides additional support for overlap in the functions of TEP-1 and CFH-1.

**Figure 4.**
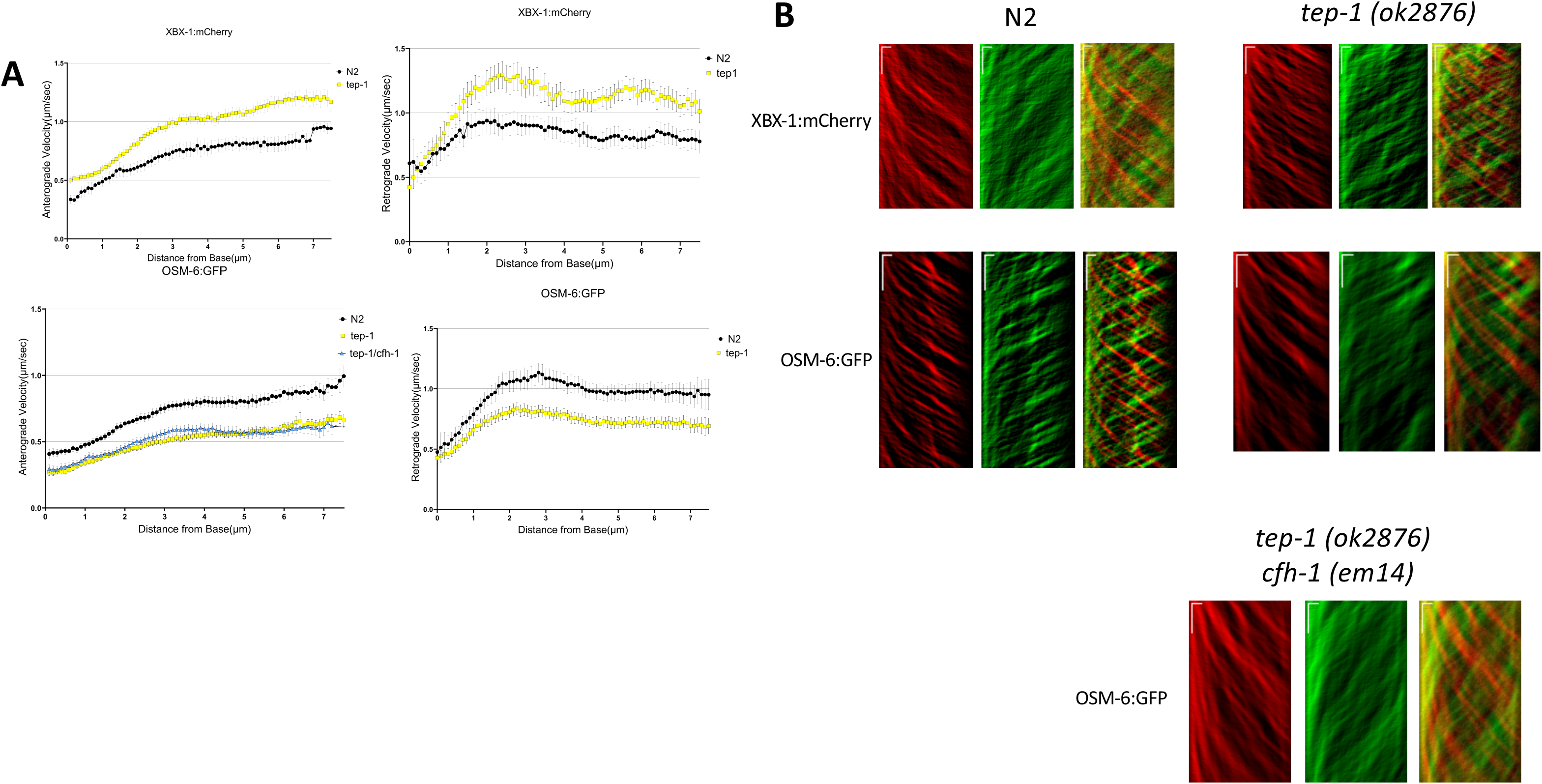
*Comparison of anterograde and retrograde movement of DYN lic /XBX-1 and IFT52/OSM-6 in WT and tep-1 mutant C. elegans day 4 adults*. A, Plots of anterograde (left) and retrograde (right) IFT velocity vs. distance from cilia base for *DYN lic /XBX-1::mCherry* (top) and IFT52/OSM-6::GFP (bottom) in phasmid sensory neurons of WT(black) and *tep-1* mutant (yellow) animals. Note that in *tep-1* mutant animals IFT velocities for *DYN lic /XBX-1* are faster than WT. In contrast, IFT velocities are decreased significantly for IFTB component, IFT52/OSM-6, in both directions in *tep-1* mutant animals. B, Representative anterograde (red) and retrograde (green) kymograph plots upon which the data are based. Horizonal scale bars represent 1μm and vertical scale bars indicate a 5 second interval.

**Figure 5.**
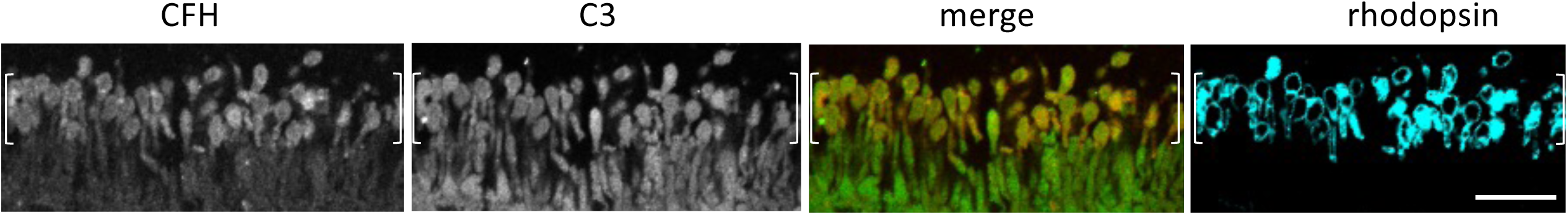
CFH and C3 localization on human photoreceptors. Antibody staining of human retina sections reveals CFH staining of photoreceptor outer segments and C3 staining of photoreceptor inner and outer segments. Also show is rhodopsin staining to indicate rod outer segments (brackets). Scale bar is 20μm.

### CFH and Thioester protein C3 colocalize on rods in human retina

CFH-1 and TEP-1 localization and phenotypes, combined with prior demonstrations of CFH localization on vertebrate photoreceptor outer segments (14,22,23,30,31), and cilia compartment defects in AMD high-risk CFH variant photoreceptors (14), raised questions regarding CFH and C3 localization in human photoreceptors. In tissue sections of human post-mortem retina, C3 is detected on both the outer and inner segments of rhodopsin positive and negative (presumably cone) cells and has significant overlap with CFH that is detected primarily on rod outer segments (identified by staining with rhodopsin) as seen previously (14,22,23,30,31).

## Discussion

The identification of CFH variants that increase disease risk (1–6) suggest that inflammation and cytolysis drive AMD pathogenesis given the well-known role of CFH in suppressing C3 activity. However, evidence showing that human CFH and C3 have functions outside of their established role in the alternative complement pathway implies that non-canonical mechanisms may also influence disease progression (12–14,22). To gain potential insight into evolutionarily ancient but previously unknown C3 functions, the localization and function of the *C. elegans* ancestral relative of the AMCOM family, TEP-1, was examined.

### Glial TEP-1 is necessary for sensory neuron inversin compartment organization

TEP-1 localizes to multiple epithelia and at the distal ends of select sensory neuron glial cells, including CEPsh, Phso, and Amso. Like CFH-1, TEP-1 localization on CEPsh contributes to the organization and maintenance of inversin/NPHP-2 compartment boundaries in sensory neurons (14) (Fig. 3). Mutations in *tep-1* and *cfh-1* mutant animals have little or no detectable effect on the distribution of the MS component ARL-13::mNG in CEP or phasmid sensory neurons, indicating that these proteins function with specificity for inversin/NPHP-2 and are not required for global cilia structure and organization.

The *tep-1(ok2874)* inversin compartment defect is effectively rescued by TEP-1 with a GPI anchor, when expressed under the control of a glial promoter but not under the control of a neuronal promoter (Fig.3), indicating that a glial membrane-associated protein maintains cilia compartment organization in an adjacent neuron. Cross-talk between glia and neurons regulate neuronal outgrowth, shape, synaptogenesis, and function, and this study strengthens links between a glial cell-surface protein and cilia organization and function in ensheathed sensory neurons (17,32,33).

In vertebrate retina, C3 is produced by Müller glia that provide support to local photoreceptors (34,35). C3 is also produced by retinal pigment epithelial (RPE) cells that while not technically glial, have a glial-like interdependent relationship with photoreceptors and engulf shed discs from photoreceptor distal ends that are embedded in their apical surface (22,34–36).

In *C. elegans*, glial socket cells ensheath the distal ends of sensory neuron cilia, suggesting the possibilities of similar interdependent relationships and functional parallels with RPE cells (17,32).

### Tep-1 modulates intraflagellar transport

Although the established roles of AMCOM family members are primarily in the innate and adaptive immune systems, the presence of an AMCOM family member in *C. elegans* suggests that there may be ancient functions for these proteins that predate separation of the nematode and vertebrate lineages. Data unexpectedly demonstrating a protective role for C3 on photoreceptors in mice may be indicative of a previously unknown function (22).

TEP-1 localization in close proximity to phasmid cilia coupled with the overlapping roles of CFH-1 and TEP-1 in inversin compartment organization (Fig. 3), and the observation that IFTB1 components behave very differently in WT and *cfh-1* mutant animals raised questions over whether TEP-1 has a similar role in regulating IFT. The finding that IFTB1 components IFT52/OSM-6 and IFT88/OSM-5 also behave very differently in WT and *tep-1* mutant animals suggests that there may be some heterogeneity in IFTB1 composition in the absence of TEP-1. Interestingly, IFT52/OSM-6 assembly into retrograde IFT trains is delayed compared to other IFT train components in WT animals, leading to the suggestion that IFT52/OSM-6 may be post-translationally modified before addition to retrograde trains (37). One possibility is CFH-1 and TEP-1 facilitate a putative post-translational modification that promotes assembly of the IFTB1 complex.

Some of the earliest signs of AMD are thinning of the photoreceptor segment and loss of interdigitation zone (IZ) integrity in optical coherence tomography (38, 39). Although it has been suggested that photoreceptor segment thinning may be a result of a defect in cholesterol recycling, it may be worth considering that a regulatory defect in photoreceptor IFT may cause an imbalance between photoreceptor outer segment synthesis and shedding by photoreceptors, and the absorption of shed material by RPE cells at the IZ. Interestingly, mutations in the genes that produce IFT-B1 components IFT88/OSM-5 and IFT52/OSM-6 result in shortened axonemes in *C. elegans* sensory neurons, so it seems plausible that defective regulation of IFT in photoreceptors might produce a similar shortening of the photoreceptor axoneme, thinning of the photoreceptor segment, and disruption of IZ integrity in the retina (27, 38, 39).

### CFH-1 and TEP-1 have overlapping and independent functions

The inversin compartment defect in *tep-1(ok2874)* mutant animals is slightly amplified in *tep-1(ok2874)*;*cfh-1(em14)* double mutants, suggesting partial mechanistic overlap between the two genes in maintaining CEP neuron inversin compartment boundaries. This might be expected given their similar distribution, their shared specificity for inversin/NPHP-2 rather than ARL-13, and the known regulatory interaction between C3 and CFH in vertebrates. However, the observation that the effect on inversin/NPHP-2 is evident in day 1 adult *tep-1* mutants (Fig. S2) but only apparent in *cfh-1* mutants after day 2 of adulthood suggests that TEP-1 may act earlier in the organization of the inversin compartment, while CFH-1 functions primarily in maintenance of inversin compartment boundaries in aging animals (14). Moreover, in contrast to their arrangement on CEP sensilla, TEP-1 and CFH-1 associate with distinct phasmid glia. Despite this non-overlapping localization, both CFH-1 and TEP-1 have roles in inversin compartment organization and IFT in phasmid cilia, suggesting that there is mechanistic overlap between these proteins that may be indirect in phasmid sensilla.

Interestingly, established AMD models suggest that C3 should have a detrimental effect on photoreceptor health in the absence of CFH-mediated inhibition. However, knockout mouse studies suggest that C3 actually has a protective effect on photoreceptor health that is independent of its interaction with CFH (22). The data presented here support the concept that outside of their established interaction in the alternative complement pathway, CFH and C3 may have distinct and evolutionarily ancient functions that make a positive contribution to the health of photoreceptors and other sensory neurons.

## Material and Methods

### C. elegans Strains and microscopy

A list of the *C. elegans* strains used in this work is included in Table S1.

Adult C. elegans strains were immobilized for 10 minutes with 15 μm levamisole (LKT laboratories) in m9 buffer and placed on a 5% agar pad on a microscope slide. Imaging was performed with a Nikon A1R confocal microscope with Nikon Elements software.

### TEP-1 deletion and mNeon Green knock-in

CRISPR-Cas9 technology was used to delete 8 nt of genomic sequence from the *C. elegans tep-1* gene (ZK337.1) gene to create a frameshift in *tep-1* coding sequence and non-sense codon and PciI restriction site. CRISPR-Cas9 technology was also used to insert mNeon Green in-frame into *tep-1* coding sequence. The sequence of the crRNA was ACAGAGTACAAATGCAGCTG (Integrated DNA technologies). The repair template used to create the deletion had a 35nt overlap with endogenous sequence on both sides of the Cas9 cleavage site and had the sequence, GGCAAATACATGGCGTCATCGGACAGAGTACAAAT CGAACTTACATTATC GCAGCTGTGGTGTCAACAACCGCGGCGCCAGTTAA (Integrated DNA Technologies). The forward and reverse primers used to amplify mNG also had 35nt overlaps with endogenous sequence on either side of the Cas9 cleavage site, GGCAAATACATGGCGTCATCGGACAGAGTACAAATGGATCCGCCGGATCCGCCGC and TTAACTGGCGCCGCGGTTGTTGACACCACAGCTGCGAACTCTCCGGATCCGGCGG. Injection mixtures were prepared as described and injected into the gonad of N2 hermaphrodites (40).

### VAB-9::wrmScarlet construction

VAB-9::V5::wrmScarlet knock-in allele *vab-9*(syb7720) was generated using CRISPR-Cas9 technology by SunyBiotech (Fujian, China). The sequence of the crRNAs were CGGTCGGAGAGCTCTGGTGG and AGTTCAAAAAATTAGACTAT. The repair template consisted of a 485bp 5’ homology arm and a 738bp 3’ homology arm flanking the insertion site. Injections mixtures were prepared and injected into the gonads of N2 hermaphrodites. Sequencing primers were TTGCGACAATGATTTCTGGA, TCTCTTTCCCAACCTAATCC, and GGTTTTCATGTTACTGACAG.

#### Timelapse Live imaging, kymograph generation and analysis

##### Sample Preparation

To synchronize the ages of C. elegans, hermaphrodites at the L4 stage were transferred to fresh nematode growth medium (NGM) plates seeded with E. coli OP50. After two days incubation, adult worms were transferred to new plates on adult day 2 to separate them from younger worms, ensuring the collection of synchronized adult day 4 worms. On adult day 4, the worms were anesthetized and immobilized in 3 µl of 10 mM levamisole in M9 buffer on a 5% agar pad placed on a microscope slide, which was covered with a glass coverslip.

##### Imaging Setup

Phasmid cilia were imaged using a Nikon A1R confocal and super-resolution system equipped with a 100X objective lens with a Nipkow spinning disk. Time-lapse images were acquired with an exposure time of 200 ms for 30 seconds. A total of 150 images were collected and stacked for further analysis.

##### Kymograph Generation and Analysis

The stacked images were processed in ImageJ using the macro tool Kymograph Clear 2.0 (41) to generate kymographs. Kymograph Clear 2.0 automatically distinguishes moving particles based on their direction and generates kymographs for each set of directional tracks, as described previously (38). Particle tracks on the kymographs were manually traced. The generated kymographs were further analyzed with Kymograph Direct (41). The velocities of the traced particles in each direction were measured at positions along the cilium. Velocity values were compiled at 0.1 µm intervals from the ciliary base and averaged for downstream analysis.

#### Immunohistochemistry of Human retinal sections

Postmortem human eyes were embedded in paraffin and sectioned as described (30). Sectioned eyes were deparaffinized in Xylene and hydrated by washing in 100%, 95%, 70%, 50%, 30% Ethanol and H2O (5 mins each). After washing in Tris-buffered Saline with 0.1% tween for 5 mins and PBS 3 times for 5 mins, sections were blocked in 10% Donkey serum in PBS with 0.5% Triton-X-100 for 1 hour. Sections were incubated at 4°C overnight with the following primary antibodies in blocking solution: mouse monoclonal antibody to Complement factor H (1:50, custom antibody described above or Invitrogen C18/3), mouse monoclonal antibody to rhodopsin (4D2) (1:400, NBP2-59690, Novus Biologicals) polyclonal antibody to rhodopsin (1:200, PA5-85608, Invitrogen), and goat polyclonal antibody to Complement 3 (1:500, PA1-29715, Invitrogen). After washing in PBS 3 times for 10mins each, sections were incubated with the following secondary antibodies for 1 hour in RT: Donkey anti-mouse Alexa Fluor 647 conjugated secondary antibody (1:200, A31571, Thermo Fisher Scientific),Donkey anti-rabbit Alexa Fluor 488 conjugated secondary antibody (1:200, A21206, Thermo Fisher Scientific), and Donkey anti-goat Alexa Fluor™ 555 secondary antibody (1:200, A21432, THermo Fisher Scientific). After a rinse and 3 washes in PBS for 10 mins each, sections were counterstained by 5µM DAPI for 7min and washed in PBS 3 times for 20mins. Sections were mounted in Slow Fade Gold (S36936, Invitrogen by Thermo Fisher Scientific) or KPL mounting Medium (5570-0005, seracare) and sealed by cover grass and nail polish. Slides were imaged using a Nikon A1R confocal microscope with Nikon Elements software.

### Statistical analysis

One-way ANOVA followed by Tukey’s or Dunnett’s multiple comparisons test and correlation analysis were performed using GraphPad Prism version 10.2.3 for macOS (GraphPad Software, San Diego, California USA, www.graphpad.com).

## Acknowledgments

The authors thank Kevin O’Connell, Sean Wallace, Shai Shaham, and Max Heiman for their exceptional generosity in sharing *C. elegans* strains and constructs. We also thank Kevin Rossomando and Qiang Lin for technical assistance and Ashley Frakes for comments on the manuscript. J.A.M is supported by the Canadian Institutes of Health Research. Some *C. elegans* strains were provided by the CGC, which is funded by NIH Office of Research Infrastructure Programs (P40 OD010440) and the C. elegans Reverse Genetics Core Facility at the University of British Columbia, which is part of the international C. elegans Gene Knockout Consortium. Research reported in this publication was supported by the National Eye Institute of the National Institutes of Health under Award Number R01EY032868 to B.E.V. The content is solely the responsibility of the authors and does not necessarily represent the official views of the National Institutes of Health.

---

**Figure S1.**
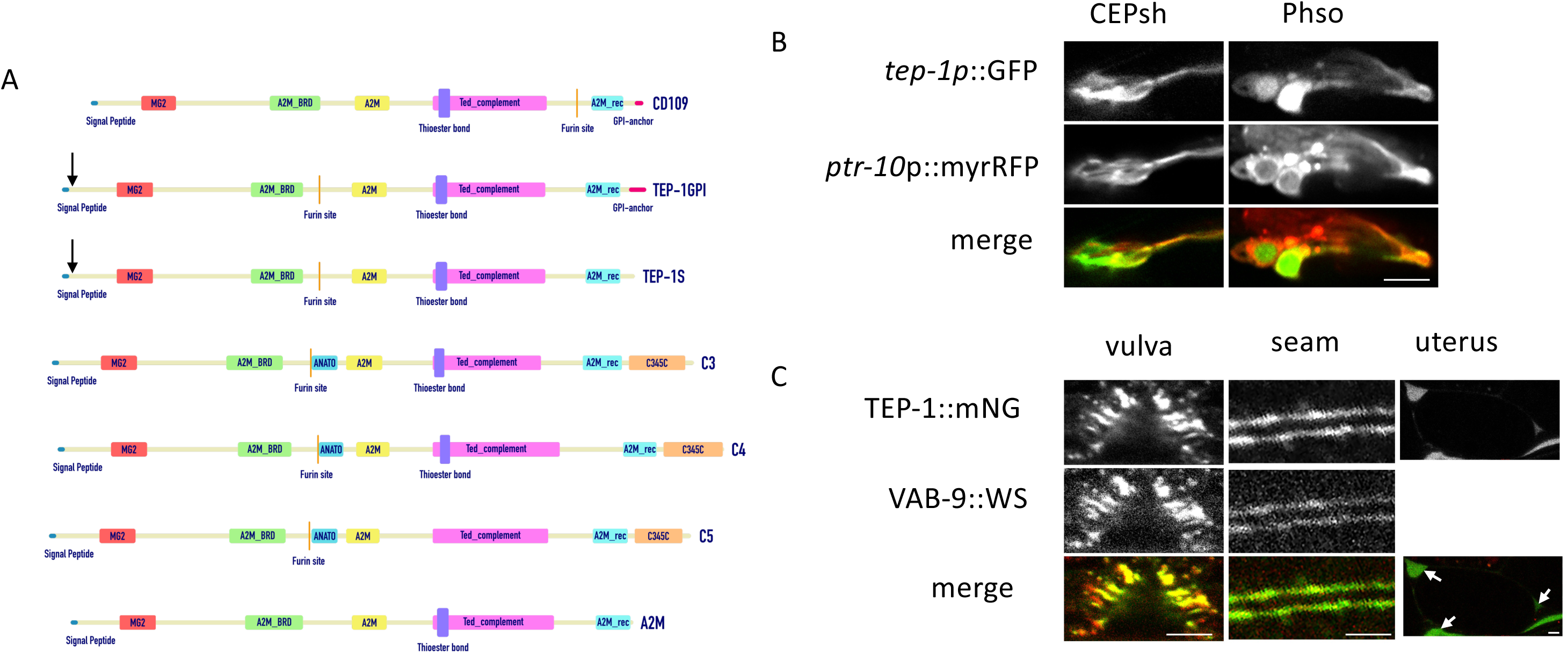
A, Comparison of the modular structures of *TEP-1 splice variant secreted form (TEP-1S) and GPI anchored form (TEP-1GPI), CD109, C3, C4, C5 and alpha-2-macroglobulin (based on 21).* Conserved domain information was compiled from <u>Interpro</u>, <u>SMART domain architecture analysis</u>, and <u>NCBI Web CD-Batch Search Tool</u>. Abbreviations are as follows: MG2:Macroglobulin domain, A2M_BRD: Alpha-2-macroglobulin bait region domain, A2M: Alpha-2-macroglobulin family, ANATO: Anaphylatoxin homologous domain, Ted_complement: complement thioester domain, A2M_rec: Alpha-2-macroglobulin receptor domain, C345C: NTR (netrin) module. Arrow shows location of CRISPR mediated insertion of mNeon Green. B, *tep-1 expression by CEP sheath and phasmid socket glia.* GFP expressed under the control of *tep-1* 5’ untranslated regulatory sequence overlaps with myrRFP under the control of ptr-10 5’ regulatory sequences in CEPsh and Phso glial cells in young adult animal. Anterior is oriented left. Scale bars is 5μm. C, Confocal image of TEP-1::mNG translational fusion in tandem with VAB-9::wrmScarlet in vulval and seam epithelial cells and TEP-1::mNG in uterine fluid surrounding a fertilized egg (arrows). Scale bar is 5μm. Dorsal is oriented up.

**Figure S2.**
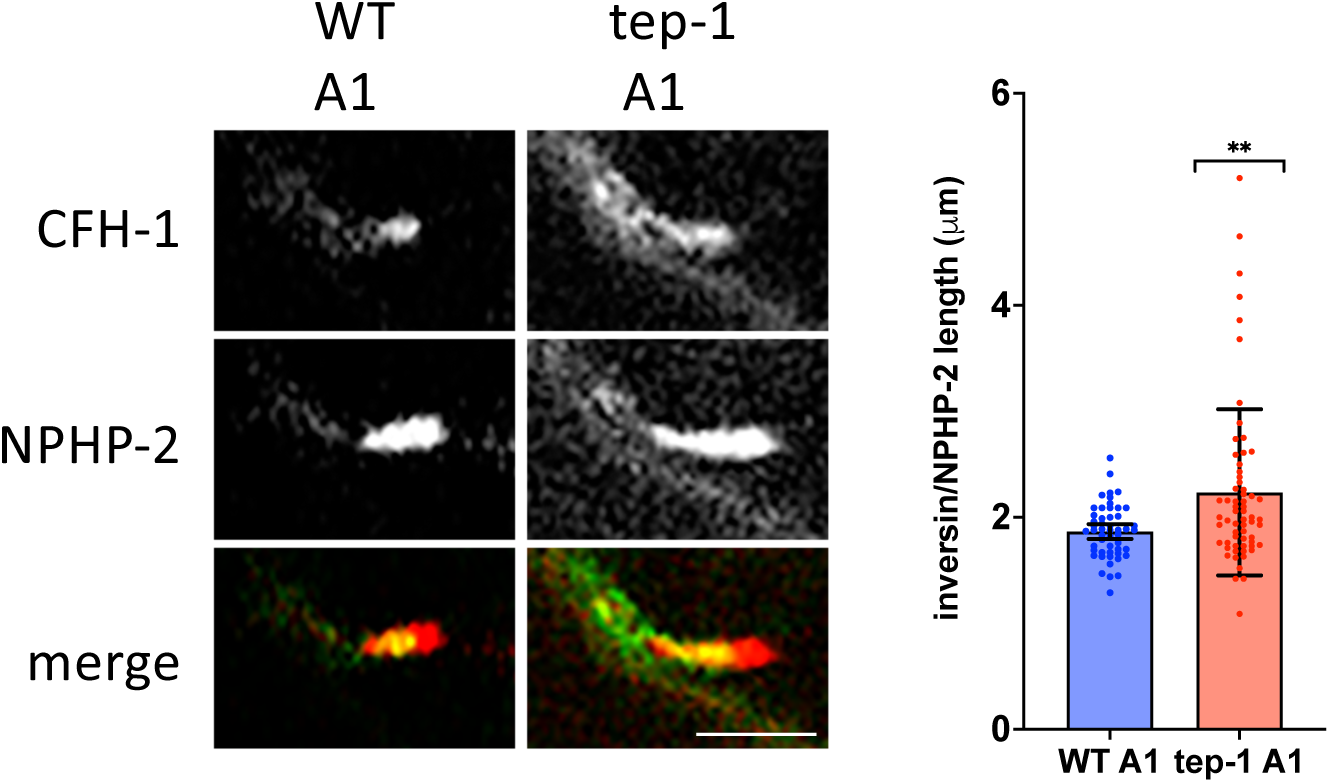
A, Comparison of CFH-1::GFP and inversin/NPHP-2::mCherry localization in CEP neurons of WT and *tep-1* mutant day 1 adults. Right, scatter plots of inversin/NPHP-2 length in CEP neurons of day 1 adult animals. Error bars indicate 95% confidence intervals. Significance is indicated by brackets with asterisks as follows: ** P<.01. Scale bar is 2.5μm.

**Figure S3.**
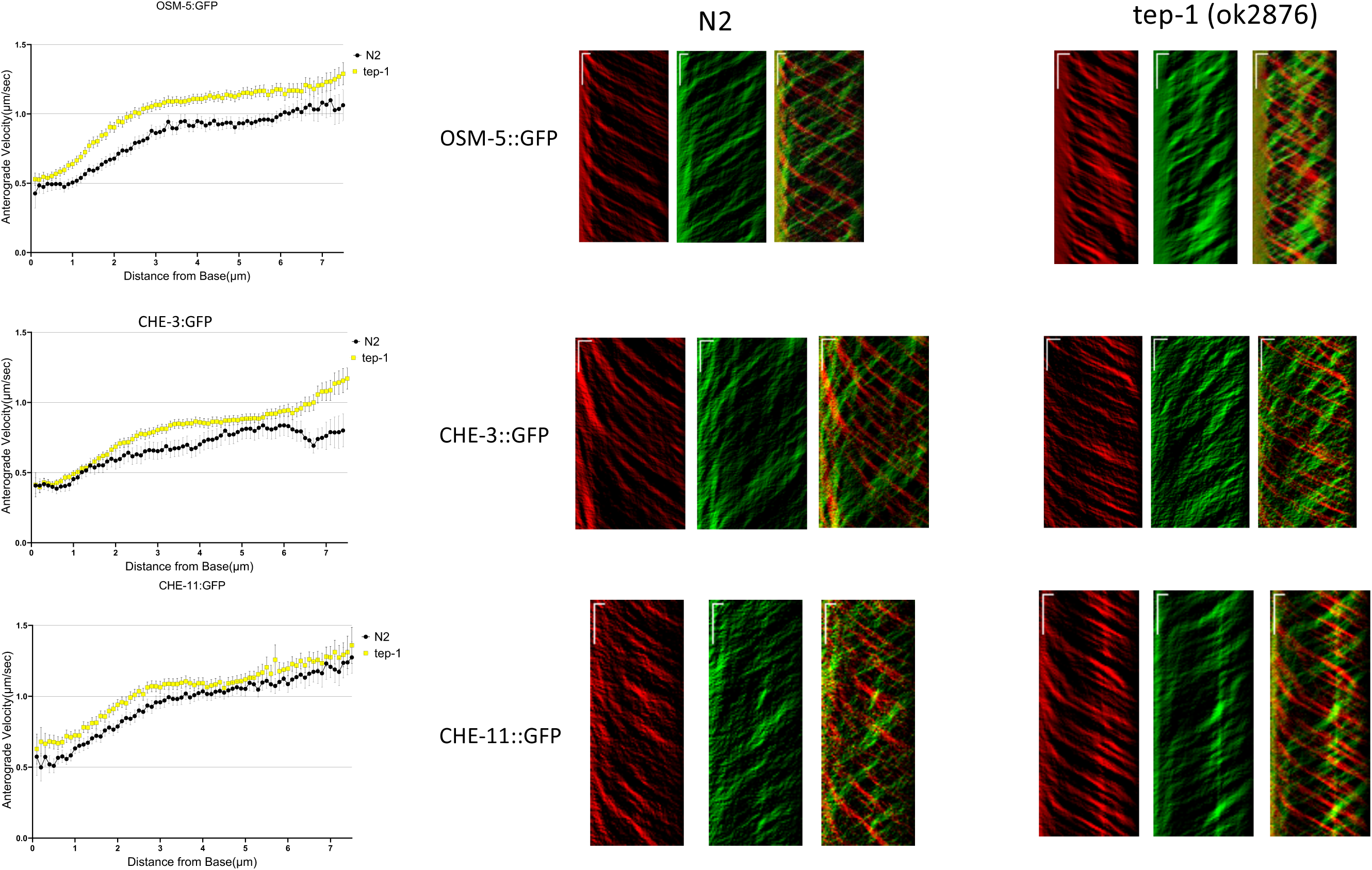
*Comparison of anterograde movement of IFT88/OSM-5,* dynein heavy chain CHE-3, and CHE-11/IFT140 in *WT and tep-1 mutant C. elegans day 4 adults*. A, Plots of anterograde IFT velocity vs. distance from cilia base for IFT88/OSM-5::GFP (top), dynein heavy chain CHE-3, and CHE-11/IFT140 (bottom) in phasmid sensory neurons of WT(black) and *tep-1* mutant (yellow) animals. Note that in *tep-1* mutant animals IFT velocities are slightly faster than WT in anterograde IFT. B, Representative anterograde (red) and retrograde (green) kymograph plots upon which the data are based. Horizonal scale bars represent 1μm and vertical scale bars indicate a 5 second interval.

**Supplemental Table S1.** List of strains used.

| STRAIN | GENOTYPE |
| --- | --- |
| BT40 | *em16* [CFH-1::eGFP]; *emls5* [dat-1p::NPHP-2::mCherry+rol-6(su1006)] |
| BT41 | *emls5* [dat-1p::NPHP-2::mCherry+rol-6(su1006)]; *oq118* [ARL-13::mNG] |
| BT42 | *cfh-1*(*em14*); *emls5* [dat-1p::NPHP-2::mCherry+rol-6(su1006)]; *oq118* [ARL-13::mNG] |
|  | *myEx1*[*Posm-5*::OSM-5::GFP + *pRF4*]; *xbx-1*(*cas502*[*xbx-1*::tagRFP]) |
| BT62 | *tep-1*(*ok2874*); *myEx1*[*Posm-5*::OSM-5::GFP + *pRF4*]; *xbx-1*(*cas502*[*xbx-1*::tagRFP]) |
| BT66 | *myEx10*[*Pche-11*::CHE-11::GFP + *pRF4*]; *xbx-1*(*cas502*[*xbx-1*::tagRFP]) |
| BT72 | *dpy-5*(*e907*); *sEX14690*[*tep-1*p::GFP+ *dpy-5*]; *ns108*[*ptr-10*p::myrRFP] |
| BT73 | *em19*[ *TEP-1*::mNG]; *ns108*[*ptr-10*p::myrRFP] |
| BT74 | *em19*[ *TEP-1*::mNG]; *emEX*[*grl-2*p::mApple+rol-6(su1006)] |
| BT75 | *em19*[ *TEP-1*::mNG]; *syb7720* [VAB-9::wormScarlet] |
| BT76 | *em19*[ *TEP-1*::mNG]; *emEX*[*bbs-8*p::XBX-1::tag::RFP] |
| BT77 | *em19*[ *TEP-1*::mNG]; *emls5*[*dat-1*p::NPHP-2::mCherry+rol-6(su1006)] |
| BT78 | *em19*[ *TEP-1*::mNG]; *emEX*[*bbs-8*p::NPHP-2::mCherry+rol-6(su1006)] |
| BT79 | *em16*[ *CFH-1*::eGFP]; *em20*[*TEP-1*::wormScarlet] |
| BT80 | *em16*[ *CFH-1*::eGFP]; *emEX*[*bbs-8*p::NPHP-2::mCherry+rol-6(su1006)] |
| BT81 | *tep-1*(*em18*); *em16* [CFH-1::eGFP]; *emls5* [dat-1p::NPHP-2::mCherry+rol-6(su1006)] |
| BT82 | *tep-1*(*ok2874*); *em16* [CFH-1::eGFP]; *emls5* [dat-1p::NPHP-2::mCherry+rol-6(su1006)] |
| BT83 | *cfh-1*(*em14*); *tep-1*(*ok2874*); *emls5* [dat-1p::NPHP-2::mCherry+rol-6(su1006)] |
| BT84 | *tep-1*(*ok2874*); *em16* [CFH-1::eGFP]; *emls5* [dat-1p::NPHP-2::mCherry+rol-6(su1006)] +*emEX*[*tep-1*p::*tep-1*+unc-122p::RFP] |
| BT85 | *tep-1*(*ok2874*); *em16* [CFH-1::eGFP]; *emls5* [dat-1p::NPHP-2::mCherry+rol-6(su1006)] +*emEX*[*ptr-10*p::*tep-1*gpi+unc-122p::RFP] |
| BT86 | *tep-1(ok2874); em16 [CFH-1::eGFP]; emls5 [dat-1p::NPHP-2::mCherry+rol-6(su1006)] +emEX[ptr-10p::tep-1s+unc-122p::RFP]* |
| BT87 | *tep-1(ok2874); em16 [CFH-1::eGFP]; emls5 [dat-1p::NPHP-2::mCherry+rol-6(su1006)] +emEX[bbs-8p::tep-1gpi+unc-122p::RFP]* |
| BT88 | *tep-1(ok2874); em16 [CFH-1::eGFP]; emls5 [dat-1p::NPHP-2::mCherry+rol-6(su1006)] +emEX[bbs-8p::tep-1s+unc-122p::RFP]* |
| BT89 | *emEX[bbs-8p::NPHP-2::mCherry+rol-6(su1006)]* |
| BT90 | *tep-1(ok2874); emEX[bbs-8p::NPHP-2::mCherry+rol-6(su1006)]* |
| BT91 | *cfh-1(em14); emEX[bbs-8p::NPHP-2::mCherry+rol-6(su1006)]* |
| BT92 | *cfh-1(em14); tep-1(ok2874); emEX[bbs-8p::NPHP-2::mCherry+rol-6(su1006)]* |
| BT93 | *tep-1(ok2874); emls5 [dat-1p::NPHP-2::mCherry+rol-6(su1006)]; oq118 [ARL-13::mNG]* |
| BT94 | *tep-1(ok2874); che-3(cas443[gfp::che-3]); xbx-1(cas502[xbx-1::tagRFP])* |
| BT95 | *tep-1(ok2874); myEx10[Pche-11::CHE-11::GFP + pRF4]; xbx-1(cas502[xbx-1::tagRFP])* |
| GOU2162 | *che-3(cas443[GFP::che-3]); xbx-1(cas502[xbx-1::tagRFP])* |

